# Metrics Matter: Why We Need to Stop Using Silhouette in Single-Cell Benchmarking

**DOI:** 10.1101/2025.01.21.634098

**Authors:** Pia Rautenstrauch, Uwe Ohler

## Abstract

Current-day single-cell studies comprise complex data sets affected by nested batch effects caused by technical and biological factors, relying on advanced integration methods. Silhouette is an established metric for assessing clustering results, comparing within-cluster cohesion to between-cluster separation, and adaptations of it have emerged as the dominant choice to evaluate the success of these integration methods. However, silhouette’s assumptions are often violated in single-cell data integration scenarios. We demonstrate that silhouette-based metrics can neither reliably assess batch effect removal nor biological signal conservation and are thus inherently unsuitable for data with (nested) batch effects. We propose alternative, robust evaluation strategies that enable accurate integration method assessment and call to update benchmarking practices.

## Main text

Integrating single-cell data remains a key challenge of single-cell analysis due to the increasing complexity and volume of data sets generated. These data sets often include intricate, nested batch effects from both technical and biological factors, requiring rigorous evaluation of integration methods to ensure accurate integration and interpretation. Silhouette-based evaluation metrics have become widely adopted to address this challenge. As an integral part of current data integration benchmarking, they are used for scoring both biological signal conservation (bio-conservation) and batch removal. However, we demonstrate that these metrics cannot reliably score data integration.

The metric “silhouette” scores clustering quality by comparing within-cluster cohesion to between-cluster separation (Rousseeuw, 1987), and was originally developed for evaluating unsupervised clustering results of unlabeled data (internal evaluation). In the single-cell field, silhouette was thus quickly taken up for determining the optimal number of clusters in single-cell data sets (Wagner et al., 2016; Scialdone et al. 2016). More recently, silhouette has been adapted for evaluating horizontal data integration (Argelaguet et al., 2021), for instance, to score bio-conservation by assessing how well cell type annotations (based on labeled data, i.e., external evaluation) from distinct batches co-cluster (Haghverdi et al., 2018; Tran et al., 2020; Luecken et al., 2022). From 2017 onwards, silhouette-based metrics have also been employed for scoring batch effect removal, another key challenge of horizontal data integration (Risso et al., 2018; Büttner et al., 2019; Cole et al., 2019). Here, the silhouette concept is, however, inverted for scoring how well cells from distinct batches (external labels) mix. Fueled by a large-scale single-cell benchmark and accompanying toolbox, silhouette-based batch removal metrics have become a predominant score to evaluate and claim the success of many new single-cell integration methods (Luecken et al., 2021; Luecken et al., 2022).

Unfortunately, it appears to have gone unnoticed that silhouette-based batch removal metrics completely fail when scoring data integration in even modestly challenging scenarios. To illustrate this, consider a simplified, illustrative example: we simulate four single-cell RNA-seq samples with three cell types. The samples are split into two groups, mimicking that they were sequenced at two distinct sites (Figure 1(a)). This corresponds to data with batch effects nested in groups with decreasing levels of between-group batch effects (or, conversely, increasing levels of successful data integration), which we complement with an overcorrected scenario. To evaluate the behavior of silhouette scores for evaluating batch removal, we chose the ‘ASW batch’ metric, a commonly used cell-type dependent implementation of a silhouette-based batch removal metric (scib package (Luecken et al., 2022)). We find that Batch ASW results in near maximal, close to identical scores for every scenario - no matter whether data was actually integrated or not. Silhouette scores only consider the nearest neighboring clusters - here, assigned by sample - and when samples from the same group are highly similar, batch effects between the groups cannot be captured (Figure 1 (b)). Given the increasing prevalence of nested batch effects in single-cell studies, addressing this limitation is pivotal for ensuring reliable data integration. As we will see, the problem results from the underlying definition of the silhouette score, thus extending to **every** silhouette-based metric for batch removal.

**Figure 1:**
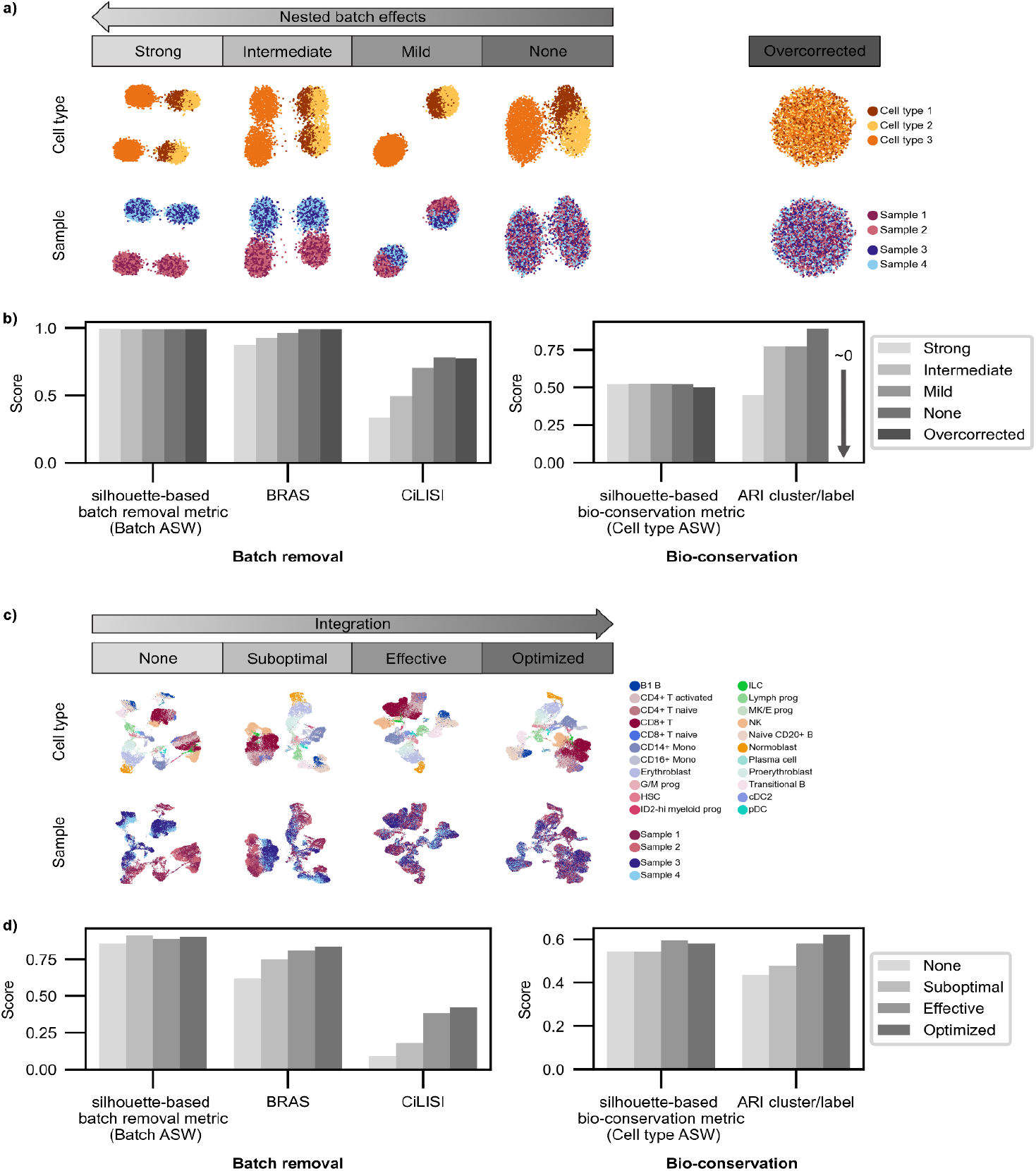
Silhouette-based metrics (Batch ASW) are unreliable with nested batch effects, failing single-cell data integration evaluation. **(a)** UMAPs of simulated data with nested batch effects between groups of samples with decreasing levels of batch effects between groups. Colored by cell type and sample. **(b)** Batch removal metrics: Unreliable metric (Batch ASW), reimplementation fixing erratic behavior called batch removal adapted silhouette (BRAS), and an alternative cell type-dependent diversity score: CiLISI. Bio-conservation metrics: Cell type ASW and ARI. **(c)** UMAPs of NeurIPS data minimal example with nested batch effects integrated with increasing success, colored by cell type and sample. **(d)** Metrics as in (b).

The silhouette is defined as follows. For a cell *i* assigned to a cluster *C*_*k*_. Given *a*_*i*_: the mean distance between a cell *i* and all other cells in the same cluster *C*_*k*_. With: *b*_*i*_: the mean distance between a cell *i* and all other cells in the **nearest** (neighboring) other cluster *C*_*l*_ where *l* ≠ *k*, the silhouette coefficient of a single cell *i*, denoted as *s*_*i*_ is given by:

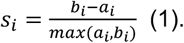

Note that this is only defined for 2 ≤ # *clusters* ≤ # *cells* − 1 and ranges between -1 and 1, with 1 indicating good cluster separation (*a*_*i*_ ≪ *b*_*i*_), values near 0 indicating cluster overlapping (*a*_*i*_ = *b*_*i*_), and -1 wrong cluster assignment (*a*_*i*_ ≫ *b*_*i*_). In contrast to the use of silhouette for internal clustering evaluation (unsupervised clustering), for scoring data integration in the single-cell field, cells are not assigned to clusters in a data-driven manner, e.g., by the result of a clustering algorithm, but by external information, such as cell type or batch labels.

To illustrate why silhouette is inadequate for evaluating batch removal, consider integrating multiple data sets (samples) with a single cell type. In this context, the aim is to score cluster overlap and not separation. Silhouette-based batch removal metrics first assign the cells of distinct samples to corresponding clusters. The assumption is that silhouette values *s*_*i*_ around 0 indicate a high level of cluster overlap and, hence, batch effect removal. However, an important detail goes unnoticed: silhouette (cf. equation (1)) considers the mean distance between a cell *i* and all other cells in only the **nearest** (neighboring) other cluster *C*_*l*_ (*b*_*i*_). A value for *s*_*i*_ around 0 is thus attainable if a given cluster overlaps with just a single other cluster and could still be very distinct from all other remaining ones. This behavior is highly problematic in the presence of nested batch effects, where samples within groups are a lot more similar to each other than between groups. If samples within groups overlap, but differences remain between samples of distinct groups, silhouette-based metrics can result in maximal scores despite remaining strong batch effects, in the worst case, even favoring suboptimal methods. In practice, data sets usually comprise a multitude of cell types. Silhouette-based batch removal scores are commonly computed per cell type label and later aggregated to account for differences in cell type composition between samples (Luecken et al., 2022). Additionally, they are transformed to range between 0 and 1, with 1 indicating best performance. The same caveats apply - in the presence of nested batch effects, maximal scores are reached even if data is insufficiently integrated.

This behavior is not limited to toy examples but, in fact, painfully obvious on real data sets. We empirically discovered this issue for ‘Batch ASW’ in the context of the NeurIPS 2021 challenge (Lance et al., 2022). The benchmark data is rich in nested batch effects of samples sequenced at different sites (intra-site differences smaller than inter-site) from bone marrow mononuclear cells (Luecken et al., 2021). Choosing a scRNA-seq subset with four batches nested into two groups (sites) for clarity, we compare metric performance on unintegrated, suboptimally integrated, effectively integrated, and optimized integrated data (Figure 1(c)). Here, the silhouette-based batch removal metric Batch ASW even favors worse solutions with stronger batch effects (Figure 1(d)), with the same observations applying to the full data set (Supplementary Figure 2(b)). While we demonstrate this behavior with scRNA-seq data, this finding generalizes to any data with nested batch effects.

Single-cell integration benchmarking is an area of active research, which has seen large-scale coordinated efforts (Tran et al., 2020; Luecken et al., 2021; Luecken et al., 2022; Hu et al., 2024; Maan et al., 2024). When first introduced, silhouette-based batch removal metrics were applied to small data sets without nested batch effects (Büttner et al., 2019), with the limitations not becoming apparent. However, given the prevalence of nested batch effects in current-day data sets, silhouette’s inability to account for nested batch effects is a real concern. It is especially problematic when they are not combined with metrics that could indicate insufficient integration, but also when evaluation results are aggregated into a single summary score that obscure possible discrepancies. Two classes of metrics should be considered to score horizontal data integration: Batch removal and bio-conservation metrics (Tran et al., 2020; Luecken et al., 2022; Maan et al., 2024). Among alternatives to silhouette, some batch removal metrics score local batch mixing and are thus not prone to the same behavior, either without cell type labels (iLISI (Korsunsky et al., 2019), kBET (Büttner et al., 2019)) or accounting for cell type imbalance if cell type labels are available (CiLISI (Andreatta et al., 2024)). Concerning bio-conservation, many clustering metrics have been applied to cell type labels (ARI, NMI, cell type ASW). Evaluating performance on a high confidence subset, e.g., samples from the same donor or technical replicates, can be a valuable option (Rautenstrauch & Ohler, 2024).

Combining local mixing batch removal with bio-conservation metrics on a cell type level has proven to be a successful strategy for evaluating integration performance (Andreatta et al., 2024). For example, applying CiLISI with ARI is robust to nested batch effects, leading to accurate rankings in our simulated and real data scenarios while flagging overcorrection (high batch removal but low bio-conservation scores) (Figure 1(b) and (d)). It is also possible to “fix” the silhouette-based metric Batch ASW to be robust to nested batch effects by redefining *b*_*i*_ as the mean distance between a cell *i* and all other cells in **any** other cluster *C*_*l*_ with *l* ≠ *k*. Changing euclidean to cosine distance results in higher discriminative power (cf. Methods for further details). This adaptation, which we call batch removal adapted silhouette (BRAS), could also be employed in other metric variants. Like CiLISI, the BRAS metric also accurately ranks simulated and real data (cf. Figure 1(b) and (d) and Supplementary Figures 1(a) and (b) and (2)).

Silhouette score problems are not limited to batch integration but also arise in scores adapted for bio-conservation. As such, the Cell type ASW score shows significant limitations in discriminating between scenarios (Figure 1(b) and (d); details concerning other bio-conservation metrics can be found in Supplementary Note 1). This limitation also goes back to repurposing the silhouette score - originally intended for internal - to external evaluation, which imposes cluster labels on the data. Highly non-convex cluster shapes, particularly in the presence of strong batch effects, cause unintended behavior as silhouette’s comparison of within-cluster cohesion to between-cluster separation becomes erratic, which can also affect batch removal metrics. Arguably, such edge cases can and have been flagged by complementing Cell type ASW with batch removal metrics (Haghverdi et al., 2018), similarly to the strategy that we show to flag the overcorrection scenario (Figure 1(b)). However, current benchmarking practices often aggregate scores across different metrics without identifying outliers. This practice can lead to misleading evaluations, as high scores from unreliable metrics can disproportionately influence the overall assessment of a method’s performance.

Single-cell data integration remains a key computational challenge and an active area of research. Our investigation reveals the inadequacy of currently prevalent silhouette-based evaluation metrics for assessing data integration. In the presence of nested batch effects, these metrics can produce near-maximal scores even when data integration fails, as they focus solely on the nearest neighboring samples. We propose a robust evaluation strategy that combines local batch mixing with bio-conservation metrics, along with modifications to the silhouette metric to address its current issues. In any case, including a baseline model, such as unintegrated data, is essential for meaningful evaluation of integration. The limitations of silhouette metrics extend to bio-conservation assessments, where non-convex cluster shapes resulting from batch effects lead to erratic behavior. In summary, silhouette-based integration metrics are inadequate and should not be used to evaluate integration. Benchmarking practices need to discontinue the use of silhouette-based metrics, especially when aggregating results.

This is required to ensure reliable assessments of integration methods, as method choice impacts downstream analyses.

## Acknowledgements

We wish to thank Michael I. Love from UNC-Chapel Hill for constructive feedback and encouragement.

## Online methods

### Data

#### Simulated data

Drawing inspiration from Andreatta et al. (2024) and a recommendation of the Splatter developer (https://github.com/Oshlack/splatter/issues/99), we simulate five scenarios with decreasing levels of nested batch effects with the Splatter package (Zappia et al., 2017) (version 1.26.0). Each scenario is composed of four samples with three cell types nested in two groups, meaning that the samples within a group are more similar to each other than between the groups. The scenarios are “Strong”, “Intermediate”, and “Mild”, as well as “None” - with no (nested) batch effects, and an “Overcorrected” scenario, with neither nested batch effects nor biological cell type signal. We first simulate data with two samples of 2000 cells stemming from three distinct cell types with varying proportions. We vary the nested batch effect for the different scenarios via the batch.facLoc and batch.facScale parameters. We then select half of the cells of the two samples, and add small noise factors to them, resulting in four samples nested into two groups of 1000 cells each. The noise factor stems from another simulated data matrix without batch and cell type structure where we use a small library size parameter lib.scale. In the “Overcorrected” scenario, we choose no differential expression between cell types and samples.

#### Real data

We employ a benchmarking data set from the NeurIPS 2021 Multimodal Single-Cell Data Integration competition, specifically designed to contain nested batch effects for evaluating integration. In particular, Luecken et al. (2021) profiled bone marrow mononuclear cells from multiple donors across distinct sites, with inter-site batch effects being larger than intra-site batch effects between donors. For demonstration purposes, we only use the scRNA-seq data of the Multiome data accessible via GEO accession: GSE194122, in particular, a preprocessed AnnData object provided as a supplementary file. We further used a minimal data subset (minimal example) to illustrate the unreliable behavior of silhouette-based metrics with nested batch effects with four samples from four donors from two distinct sites s1d1, s1d3, s4d8, and s4d9, for our main figure panels, which we renamed to Sample 1, 2, 3, and 4, respectively. We also consider the full data set, with results shown in Supplementary Figure 2.

### Data integration

#### Simulated data

No integration was performed, as we have simulated differing levels of nested batch effects, which can in turn be interpreted as varying success at batch effect removal.

#### Real data

To demonstrate the insensitivity of silhouette-based batch removal metrics to differing levels of nested batch effects, we aimed to obtain integration results with varying success. The data was first normalized to median total counts and logarithmized, and then dimensionality reduced with PCA. No integration (“None”) serves as a baseline. A naive, mild batch correction (“Suboptimal”) was achieved by batch-aware selection of highly variable genes (hvg), prioritizing genes that are highly variable across batches, which is applied before PCA (carried out with scanpy (Wolf et al., 2018)). To obtain different batch removal strengths, we used our tunable model liam (Rautenstrauch & Ohler, 2024), which gives us control over distinct batch removal strengths. In particular, we applied liam, to the raw scRNA-seq data of the BMMC Multiome data set with default parameters (“Effective”), and increased batch removal by setting the adversarial scaling parameter to 5 (“Optimized”). Of note, the findings related to the metrics are not specific to the integration models used.

### Evaluation

#### Overview

We assess horizontal data integration using a broad selection of metrics, in particular, Batch ASW, iLISI, CiLISI, BRAS, and BRAS variants for batch removal, and cLISI, Cell type ASW, NMI cluster/label and ARI cluster/label for bio-conservation.

For most metrics, we use the scib package, except for the implementations for the custom CiLISI and newly proposed BRAS metrics (detailed below).

All metrics are scaled to range between 0 and 1, with 1 being optimal. For the silhouette-based metric Cell type ASW this implies that original silhouette scores around 0 correspond to transformed scores of approximately 0.5. We use low-dimensional embeddings as input: PCA embeddings for simulated data, and PCA or liam embeddings for the NeurIPS data.

#### Custom implementations of batch removal metrics robust to nested batch effects

*CiLISI*: We implement a custom version of CiLISI (Andreatta et al., 2024), a cell-type aware version of iLISI. First, we compute iLISI (range 0-1, scib implementation) per given cell type label, which is summarized into a weighted mean (weighted by number of cells per cell type label).

*Batch removal adapted silhouette (BRAS*): To account for nested batch effects in single-cell data, we modify the silhouette score *s*_*i*_ as described in equation 1. Specifically, we redefine *b*_*i*_ as the mean distance between a cell *i* and all other cells in **any** other cluster (default in BRAS). We also test a version with *b*_*i*_ as the distance between a cell *i* and all other cells in the **furthest** other cluster.

The modified silhouette score is computed per cell *i* assigned to a cluster *C*_*k*_. Following Luecken et al.’s (2022) implementation:

*s*_*i*_ = |*s*_*i*_|, with *s*_*i*_ computed as in equation 1.

Then, for each cell type label *k* corresponding to cluster *C*_*k*_ we define the BRAS score as:

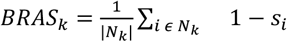

where *N*_*k*_ denotes the set of cells assigned to cluster *C*_*k*_ and |*N*_*k*_|the number of cells in that set. For the final *BRAS* score, we average over the set of unique cell labels *M*.

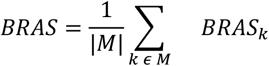

We use cosine distance as the default for BRAS, finding it provides higher discriminative power than euclidean distance (Supplementary Figure 1(a) and (b) and Supplementary Figure 2(b)). We also compute Batch ASW and Cell type ASW with cosine distance.

##### Details on ARI cluster/label and NMI cluster/label

Following Luecken et al. (2022), we optimized (Leiden) clustering with respect to the ARI and NMI metric across a range of clustering resolutions (0-2, step 0.1) and show these results in Figure 1 and Supplementary Figure 1 and 2 (Leiden is now the current default in scib, in the original publication the Louvain algorithm was used). For a discussion on potential limitations of this strategy, its impact on our results and alternative strategies see Supplementary Note 1 and Supplementary Figures 3-5.

## Code availability

The scripts and notebooks for data preprocessing, analyses, and figure generation are publicly available at https://github.com/ohlerlab/metrics_matter_manuscript_reproducibility and will be deposited in Zenodo upon acceptance.

