## Supplementary information for "Metrics Matter: Why We Need to Stop Using Silhouette in Single-Cell Benchmarking"

### Supplementary Note 1

#### Impact of clustering strategy on bio-conservation metrics ARI and NMI

To compute ARI or NMI, used here to score bio-conservation, we need to compare a clustering for any given input to ground truth labels (cell type labels). The choice of clustering algorithm and hyperparameters affects results. Luecken et al. (2022) opted to optimize clustering for the Louvain algorithm with respect to the NMI and ARI metrics across a range of clustering resolutions. This strategy can lead to cluster numbers strongly deviating from the number of ground truth cell type labels and distinct number of clusters for any given scenario, complicating comparisons and potentially favoring unrealistic solutions. Recently, Maan et al. (2024) chose to optimize clustering based on the actual number of ground truth clusters (cell types).

A recent study proved that the NMI metric can exhibit biased behavior when the number of detected clusters exceeds the true number of clusters (Mahmoudi & Jemielniak, 2024). In light of this, and due to the potential limitations of optimizing with little constraints, we sought to assess the impact of different strategies for deriving a clustering to compare to ground truth labels with ARI and NMI.

We compare the results of choosing the maximum score in the full range of tested resolutions (0-2, step 0.1) of the Leiden clustering algorithm with choosing a maximum score only for results whose number of clusters is within ±20% (bounded) of the ground truth (cell type labels). Supplementary Figures 3-5 a) show at which resolution and respective number of clusters maximal scores were reached in the full range and in the bounded region. Supplementary Figures 3-5 b) illustrate how this impacts the overall ranking of distinct scenarios for the different data sets.

We find that this choice impacts results. For example, for the full NeurIPS data, scenario “Suboptimal”, the maximum scores for ARI and NMI in the full range corresponds to a clustering output (12 clusters) that strongly deviates from the number of ground truth clusters (22) (Supplementary Figure 5 a)). In several cases, the choice of strategy even led to different rankings (e.g., ARI for NeurIPS data minimal example, Supplementary Figure 4 b)).

These findings do not affect the main conclusions of our paper regarding silhouette-based metrics, but rather underscore that exploring optimization strategies based on the number of ground truth clusters needs further investigation.

### Supplementary Figures

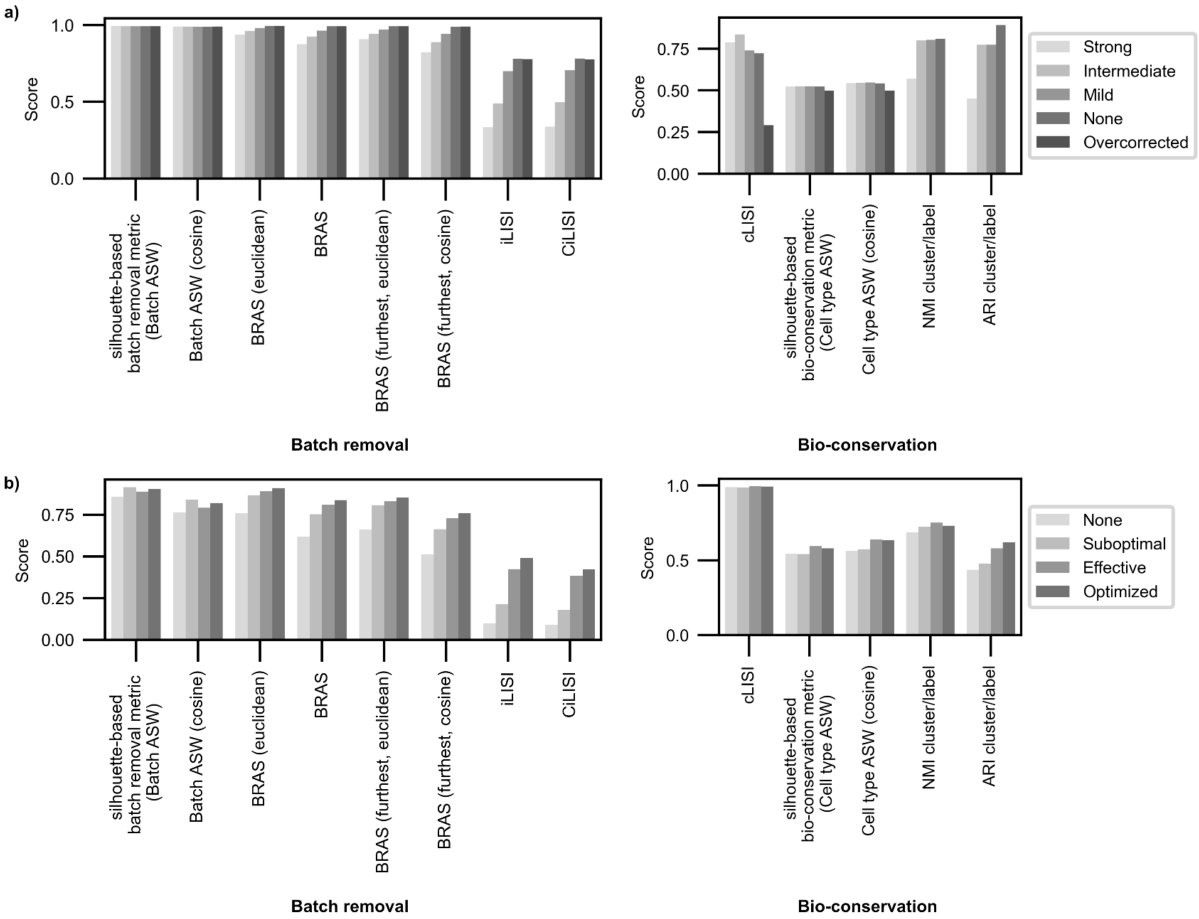


**Supplementary Figure 1:** Extended evaluation metrics. Batch removal and bio-conservation metrics **(a)** for simulated and **(b)** for real data minimal example (cf. Figure 1).

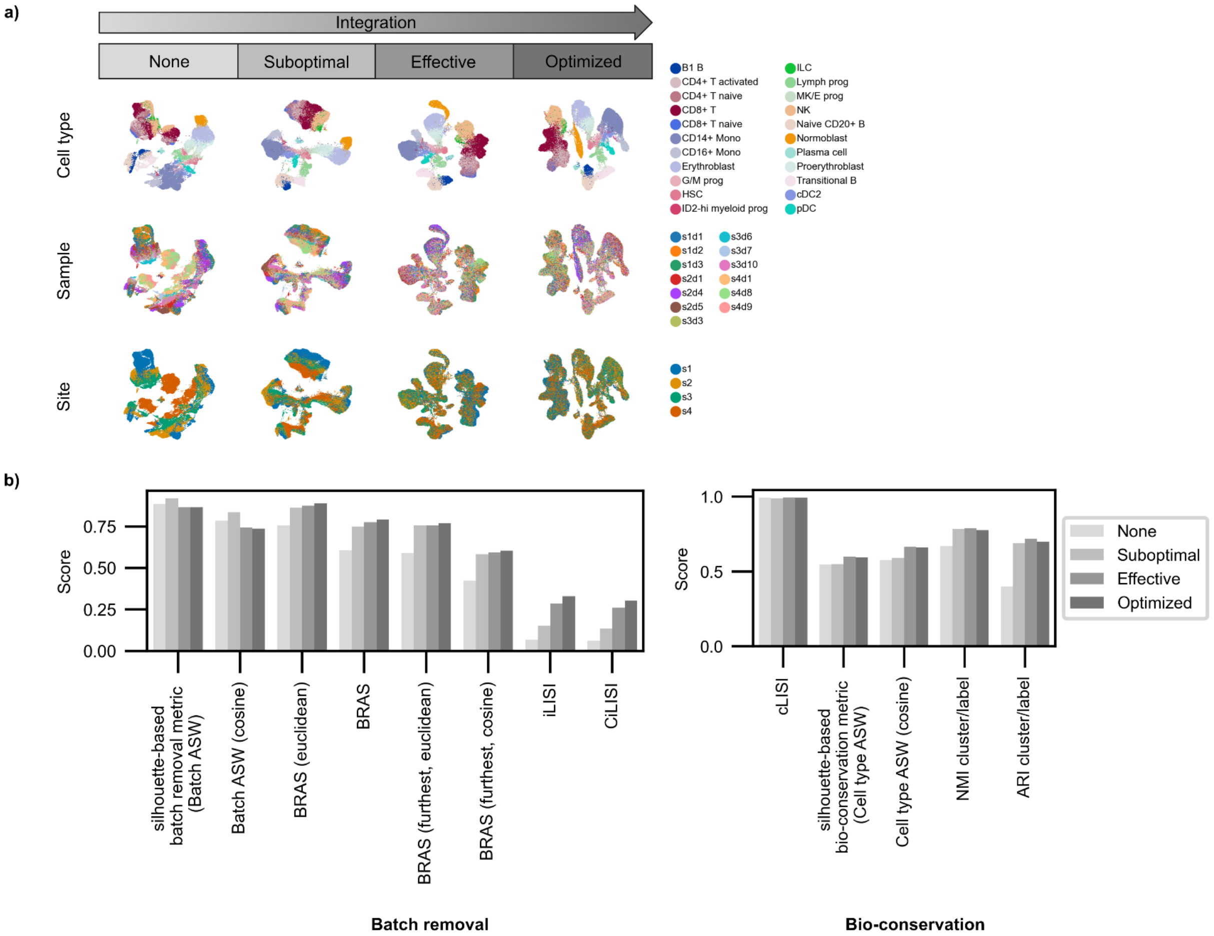


**Supplementary Figure 2: Silhouette-based metrics (Batch ASW) are unreliable with nested batch effects, failing single-cell data integration evaluation (2).**
**a)** UMAPs of full NeurIPS data set with nested batch effects integrated with increasing success, colored by cell type, sample, and site. **(b)** Extended evaluation metrics.

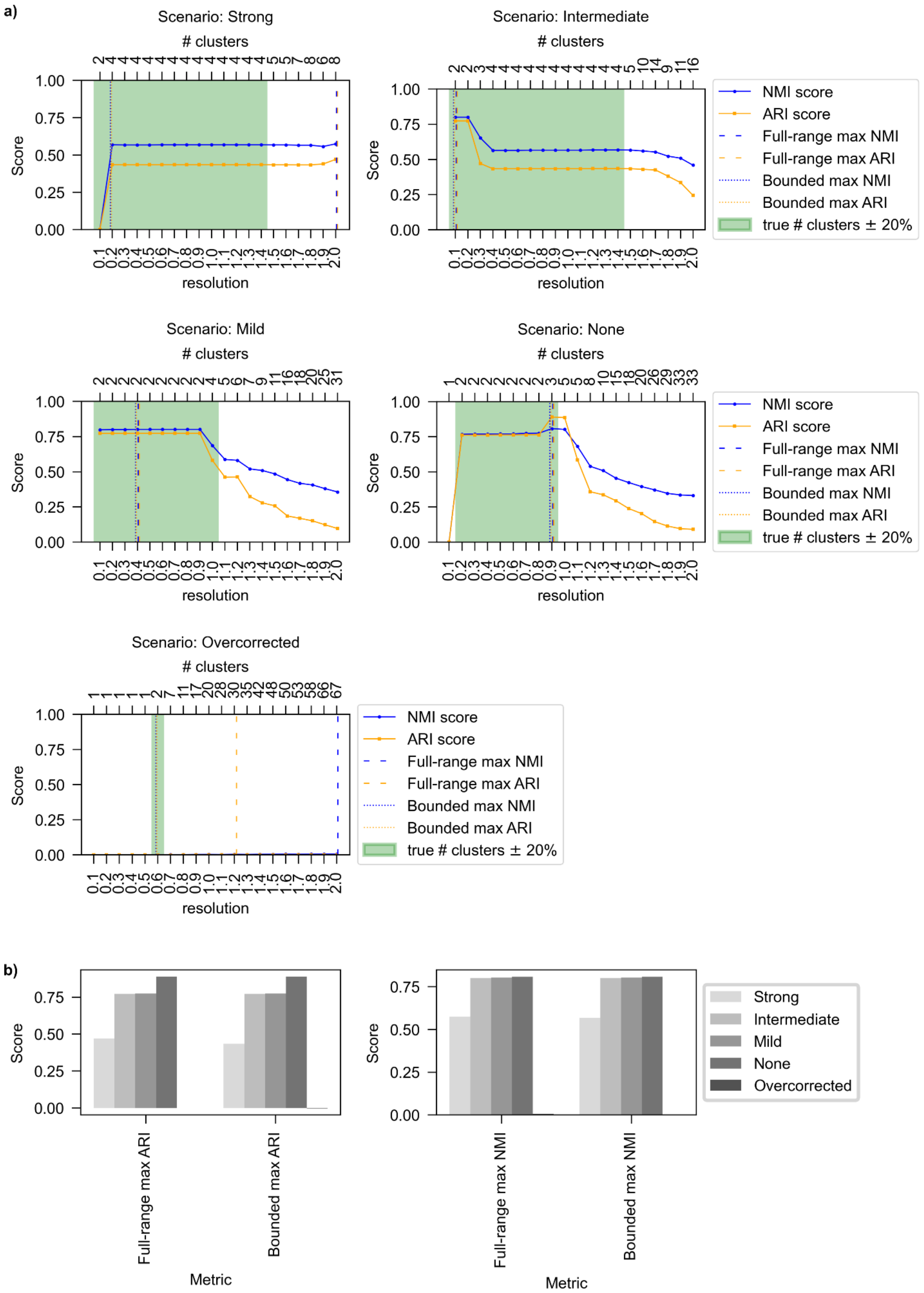


**Supplementary Figure 3: Impact of clustering strategy on ARI and NMI bio-conservation metrics for simulated data. a)** Relationship between Leiden clustering resolution (bottom x-axis), resulting cluster count (top x-axis), and corresponding ARI and NMI scores. Dashed lines indicate resolution and cluster count for maximum metric score across full range (0-2, step 0.1). Green area highlights results within ±20% of true cluster count. Dotted lines show resolution and cluster count for maximum score within bounded range. True cluster count: 3. **b)** Comparison of max scores from different clustering strategies shown in a).

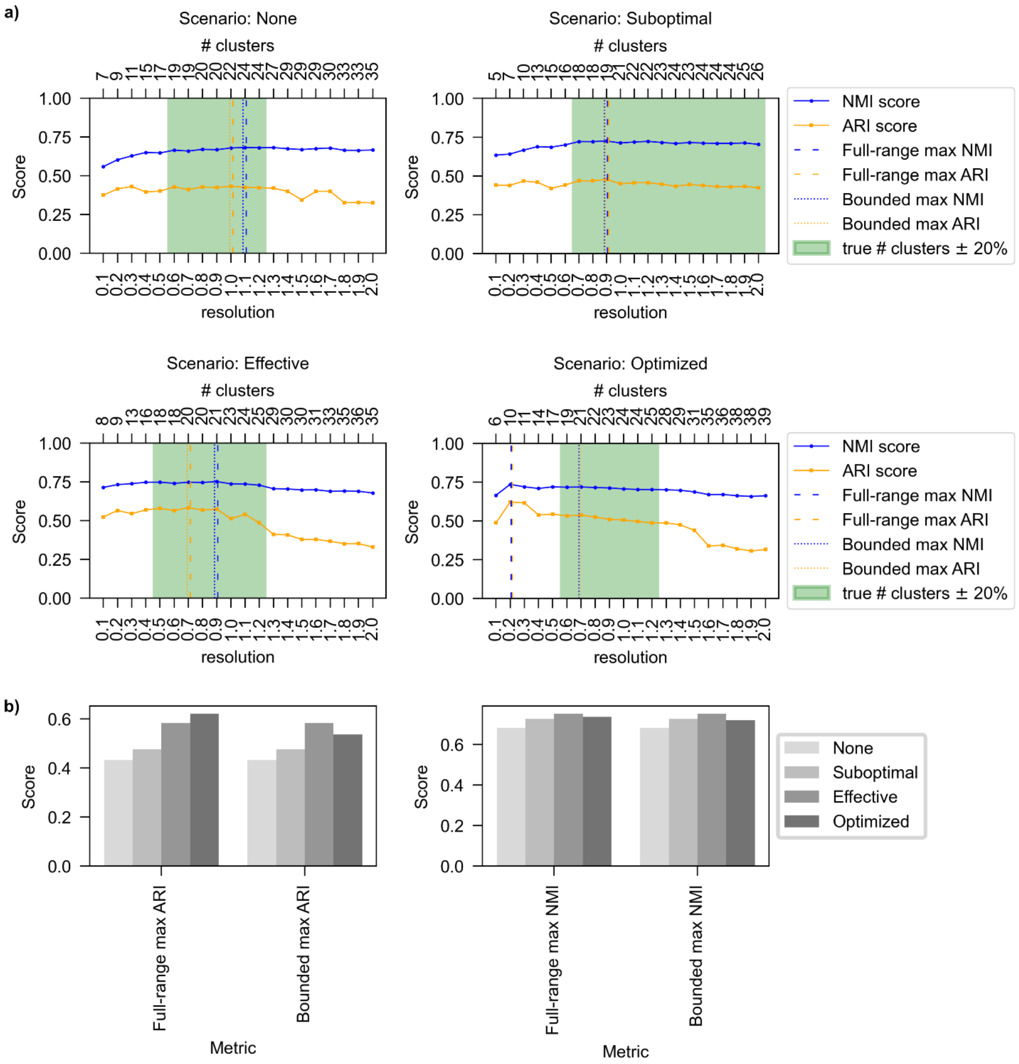


**Supplementary Figure 4: Impact of clustering strategy on ARI and NMI bio-conservation metrics for NeurIPS data minimal example. a)** Relationship between Leiden clustering resolution (bottom x-axis), resulting cluster count (top x-axis), and corresponding ARI and NMI scores. Dashed lines indicate resolution and cluster count for maximum metric score across full range (0-2, step 0.1). Green area highlights results within ±20% of true cluster count. Dotted lines show resolution and cluster count for maximum score within bounded range. True cluster count: 22. **b)** Comparison of max scores from different clustering strategies shown in a).

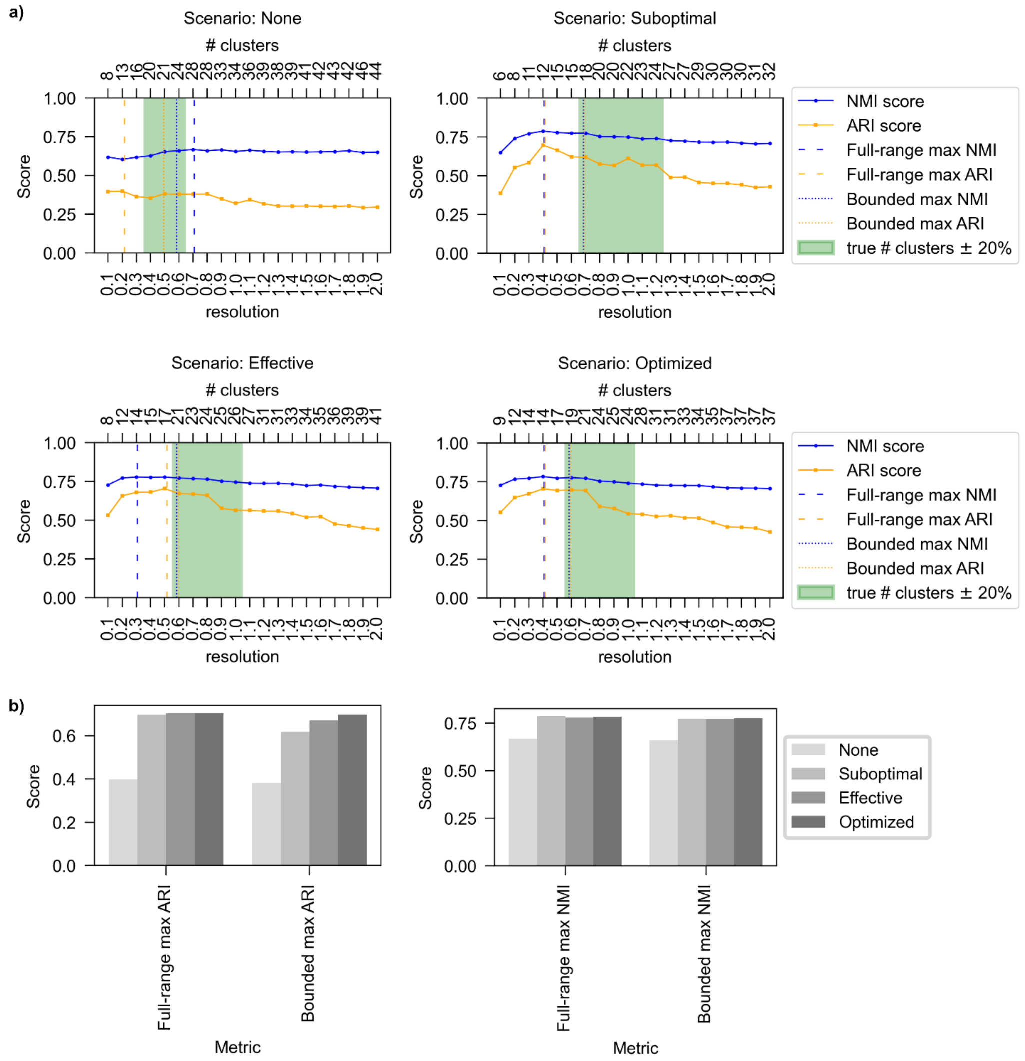


**Supplementary Figure 5: Impact of clustering strategy on ARI and NMI bio-conservation metrics for full NeurIPS data. a)** Relationship between Leiden clustering resolution (bottom x-axis), resulting cluster count (top x-axis), and corresponding ARI and NMI scores. Dashed lines indicate resolution and cluster count for maximum metric score across full range (0-2, step 0.1). Green area highlights results within ±20% of true cluster count. Dotted lines show resolution and cluster count for maximum score within bounded range. True cluster count: 22. **b)** Comparison of max scores from different clustering strategies shown in a).
